# Visibility Graph Based Community Detection for Biological Time Series

**DOI:** 10.1101/2020.03.02.973263

**Authors:** Minzhang Zheng, Sergii Domanskyi, Carlo Piermarocchi, George I. Mias

## Abstract

**Motivation:** Temporal behavior is an essential aspect of all biological systems. Time series have been previously represented as networks. Such representations must address two fundamental problems: (i) How to create the appropriate network to reflect the characteristics of biological time series. (ii) How to detect characteristic temporal patterns or events as network communities. General methods to detect communities have used metrics to compare the connectivity within a community to the connectivity one would expect in a random model, or assumed a known number of communities, or are based on the betweenness centrality of edges or nodes. However, such methods were not specifically designed for network representations of time series. We introduce a visibility-graph-based method to build networks from different kinds of biological time series and detect temporal communities within these networks.

**Results:** To characterize the uneven sampling of typical experimentally obtained biological time series, and simultaneously capture events associated to peaks and troughs, we introduce the Weighted Dual-Perspective Visibility Graph (WDPVG) for time series. To detect communities, we first find the shortest path of the network between start and end nodes to identify nodes which have high intensities. This identifies the main stem of our community detection algorithm. Then, we aggregate nodes outside the shortest path to the nodes found on the main stem based on the closest path length. Through simulation, we demonstrate the validity of our method in detecting community structures on various networks derived from simulated time series. We also confirm its effectiveness in revealing temporal communities in experimental biological time series. Our results suggest our method of visibility graph based community detection can be effective in detecting temporal biological patterns.

**Availability:** The methods of building WDPVG and visibility graph based community detection are available as a module of the open source Python package PyIOmica (https://doi.org/10.5281/zenodo.3691912) with documentation at https://pyiomica.readthedocs.io/en/latest/. The dataset and codes we used in this manuscript are publicly available at https://doi.org/10.5281/zenodo.3693984.

## 1 Introduction

Longitudinal behavior is an inherent aspect of all biological systems, and has been widely investigated in various contexts, such as systems biology (Alon, 2006), metabolic pathway analysis (Berk *et al*., 2011), and, recently, gene expression (Bar-Joseph *et al*., 2012). With the development of novel technologies in sequencing, mass spectrometry and other omics, multi-level biological time series are becoming easier to obtain. An important example is provided by longitudinal data from personal health monitoring devices. Recent studies have shown that omics time series have a wide range of applications in personal health and precision medicine. Multi-omics time series data can be used in precision health (Rose *et al*., 2019), and have provided insights into the onset of type 2 diabetes mellitus (Zhou *et al*., 2019) and lung development Ding *et al*. (2019). Omics time series can also be used to monitor health events, changes in physiological states (Piening *et al*., 2018; Stanberry *et al*., 2013) and in molecular and medical phenotypes (Chen *et al*., 2012). The rapidly increasing availability of biological time series requires new methods to integrate different types of data, analyze them, and interpret the results in a fast and informative way. Many platforms for multi-biological and multi-omics data integration have been developed, including software such as DAVID (Sherman *et al*., 2009), Galaxy (Giardine *et al*., 2005) and GenePattern (Reich *et al*., 2006), our recent frameworks MathIOmica (Mias *et al*., 2016) and PyIOmica (Domanskyi *et al*., 2019), which incorporate time-series categorization, and many more. Network-based methods have been shown to be effective in transforming time series into graph objects and capturing their characteristics, potentially allowing for faster learning approaches (Yang and Yang, 2008; Zou *et al*., 2019).

Currently several methods exist for transforming time series to complex networks. For example, complex networks have been constructed from pseudo-periodic time series by Zhang and Small (2006), who used single nodes to represent cycles, and introduced a correlation-based threshold to link node pairs. Another effective and efficient approach is to consider time series points as a series of sequential intensity bars that are then connected based their inter-visibility to obtain a *visibility graph* (VG) representation of the time series (Lacasa *et al*., 2008), which recently attracted great interdisciplinary interest (Ahmadlou *et al*., 2010; Luque *et al*., 2011; Zhu *et al*., 2014; Zou *et al*., 2014; Bhaduri and Ghosh, 2015). There are two types of VG typically considered: (i) the Natural Visibility Graph (NVG) and, (ii) the Horizontal Visibility Graph (HVG) (Luque *et al*., 2009).

To construct a VG, we consider {*s* (*t*_*x*_); *t*_*x*_ = 1, 2, 3, …, *N* } as an *N* time point series in temporal ordering. The VG is obtained by first representing the time series points as *N* nodes in a network, where nodes *i* and *j* represent times *t*_*i*_ and *t*_*j*_, with intensities *s*(*t*_*i*_) and *s*(*t*_*j*_) respectively. Edges are constructed by joining nodes *i, j* if any other intermediate time point *t*_*k*_, such as *t*_*i*_ *< t*_*k*_ *< t*_*j*_, has intensity *s*(*t*_*k*_) that satisfies the following conditions for NVG and HVG respectively:

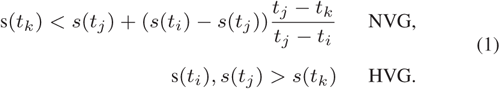

Here, in the NVG formulation, an edge is added connecting nodes *i* and *j* if any other time point *t*_*k*_, between *t*_*i*_ and *t*_*j*_, has a corresponding intensity *s*(*t*_*k*_) that lies below the line connecting *s*(*t*_*i*_) and *s*(*t*_*j*_) (i.e. there is a direct line-of-sight between these peaks). The HVG has a simpler edge construction condition: an edge is added connecting nodes *i, j* only if all intermediate intensities *s*(*t*_*k*_) are less than both *s*(*t*_*i*_) and *s*(*t*_*j*_).

NVGs and HVGs are connected networks. The VG conserves the structure of the time series in the graph topology (Lacasa *et al*., 2008). However, the HVG original constructions do not account explicitly for the potential effects of uneven time sampling or missing time points. In the NVG such uneven sampling results in different visibility of timepoints (changes in viewing angles in the construction), which implicitly incorporates the effect of time distances, but without explicitly weighting the edges the actual distance between nodes cannot be accounted for. In realistic situations outside a laboratory setting, uneven sampling occurs often. This may be due to technical limitations (for example in mass spectrometry proteomics technical replicates may still sample different proteins and lead to missing data and hence uneven sampling), or limitations in subject participation (for example in clinical trials and human subject research, the subjects’ work-dependent schedules may affect their regular participation). These shortcomings limit the traditional VG application in biological/medical time series analysis. Another limitation of the traditional VG is the inability to simultaneously capture peaks and troughs (points below the baseline). For example, the VG maps sinusoidal and cosinusoidal time series to different graphs, but these two kinds of time series should be considered equivalent up to a change of phase.

Another challenge in network representations of biological time series is the lack of specific methods for detecting temporal communities. A community is defined as a group of nodes, where the nodes within a community are tightly connected, whereas the nodes between different communities have loose connections (Girvan and Newman, 2002). Each community in complex networks representing biological time series thus identifies nodes with similar temporal behavior that are likely to represent the same biological system status. One highly effective approach for identifying communities is to compare the actual number of intracommunity edges to what one would expect by a random placement of the edges (Girvan and Newman, 2002; Clauset *et al*., 2004; Newman, 2006). This approach is based on the assumption of a random graph null model. However, VGs cannot be considered as random graphs, even if a VG is constructed based on a random time series. This is due to the sequential nature of the nodes, the resulting connected graph, and the underlying degree distribution (Luque *et al*., 2009).

In this investigation, we introduce the method of “weighted dual perspective visibility graph” (WDPVG) for mapping time series to complex networks. Our WDPVG approach considers uneven sampling effects, and simultaneously captures peaks and troughs of time series. Previously, VG edge weights have been assigned based on the arctangent of ((*s*_*j*_ − *s*_*i*_)*/*(*t*_*j*_ − *t*_*i*_)), which computes the “view angle” along the direct line-of-sight connecting one intensity peak to another (Supriya *et al*., 2016). Our method provides multiple choices for the edge weights: (i) the Euclidean distance between nodes/intensity peaks, (ii) the tangent of the view angle between two nodes, (iii) the time difference between time points corresponding to connected nodes, or (iv) none. We then combined the natural view perspective VG with the “reflected view perspective VG” introduced in the methods below to create a complex network that can capture both the positive and negative intensities changes. We note, that this is the first time that Euclidean distance has been used for edge weights in VGs, to the best of our knowledge.

We also provide a new VG community detection method, which is based on shortest path calculations between VG nodes. Our method is suitable for VGs as it does not depend on random graph null models, which are commonly used in other widely used approaches, such as Newman’s method (Newman, 2006). Briefly, as described below, we compute the shortest path of the VG between start and end nodes as a main stem. The nodes on the main stem are seeds for communities, and we then aggregate nodes outside the shortest path to their most proximal seed nodes on the main stem, where proximity is determined using graph path lengths.

We used various simulated time series to test our method effectiveness, and demonstrated that our method has high tolerance for uneven sampling and signal noise. Our comparison of our method to traditional community detection methods, such as Girvan-Newman (Girvan and Newman, 2002) and Louvain (Blondel *et al*., 2008), indicated that our approach is more suitable for VGs. To show that our method can capture biological processes we also applied it to several experimentally obtained time series, longitudinal multi-omics data from blood components in prediabetics (cytokines, glucose and haemoglobin A1c) (Zhou *et al*., 2019), saliva omics data (mean gene expression) (Domanskyi *et al*., 2019) and signals from wearable biosensors (radiation exposure) (Li *et al*., 2017).

The methods of building WDPVG and visibility graph based community detection are available as a module of the open source Python package PyIOmica(Domanskyi *et al*., 2019). The dataset and codes we used as described below are available with a Python notebook, publicly available online at https://doi.org/10.5281/zenodo.3693984.

## 2 Materials and methods

### 2.1 Weighted dual perspective visibility graph (WDPVG)

The VG can characterize time series in terms of complex network theory as it can inherit the structure properties of the time series data from which it was created. VGs are robust to noise and not affected by the selection of method parameters (e.g. cutoffs/thresholds) (Liu *et al*., 2015). VGs have been widely applied in many fields (Supriya *et al*., 2016; Stephen *et al*., 2015; Bezsudnov and Snarskii, 2014). However, as we discussed above, the VG has two disadvantages: first, it does not consider the effect of uneven sampling; second, it cannot capture the time series changes below a zero baseline. Here we provide a new method, WDPVG, to overcome these limitations that restrict VG applications to biological time series analysis.

We use the following four steps to create WDPVGs, utilizing auxiliary natural visibility graphs (NVGs).

*Step 1: NVG construction*. We create an NVG from time series {*s* (*t*_*x*_); *t*_*x*_ = 1, 2, 3, …, *N* } as it was described above by equation 1, using the NVG mapping criteria.

*Step 2: Assign edge weights between two nodes*. We have flexible choices for the edge weight between two nodes, including no weight, Euclidean distance (Equation 2), the tangent of the view angle (Equation 3), or the time difference (Equation 4):

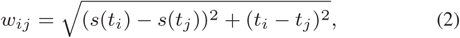

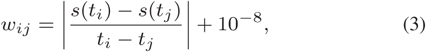

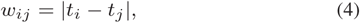

where, *w*_*ij*_ represents the weight of the edge between nodes *i* and *j*, which correspond time points *t*_*i*_ and *t*_*j*_ respectively, in time series {*s* (*t*_*x*_)}. In equation 3, we added the offset 10^−8^ to account for the case *s*(*t*_*i*_) − *s*(*t*_*j*_) = 0. The algorithm implementation details are available in the documentation of the functions in PyIOmica. In this manuscript, we use Euclidean distance between nodes as the edge weight.

After Step 2, We compute the adjacency matrix, *A*, of the normal perspective NVG.

*Step 3. Compute the reflected perspective NVG*. We invert the time series {*s* (*t*_*x*_)}, by reflecting across the time axis, i.e. for each *s*(*t*_*i*_) in {*s* (*t*_*x*_)}, let *s*^′^(*t*_*i*_) = −*s*(*t*_*i*_), where we obtain the inverted time series {*s*^′^ (*t*_*x*_)}. We then repeat Steps 1 and 2 for *s*^′^(*t*_*x*_) to get the reflected perspective NVG and the adjacency matrix *A*^′^.

*Step 4. Combine the normal perspective NVG and reflected perspective NVG*. For any pair of *i, j*, elements *A*_*ij*_ and 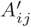 have two possible relationships: either 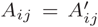, or one of them is 0 but the other one is non-zero. We can combine the *A* and *A*^′^ to get the WDPVG adjacency matrix *A*^*d*^ by the following criteria:

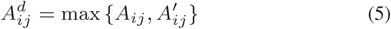

If we use the HVG mapping criteria, i.e. s(*t*_*i*_), *s*(*t*_*j*_) *> s*(*t*_*k*_) *t*_*i*_ *< t*_*k*_ *< t*_*j*_, instead of the NVG mapping criteria, we can obtain the weighted dual perspective horizontal visibility graph.

It is important to note that in case that either we are not interested in changes below the baseline, or that the intensities of the time series are all non-negative, the normal perspective weighted VG is enough, and we do not need to create a WDPVG.

### 2.2 Shortest path based community detection

The shortest path in a VG between the start node (corresponding to first time point) and end node (corresponding to last time point) identifies a bundle of nodes which have high intensities, and thus is the determining factor for the entire network structure. This shortest path acts as a stem for community identification in VG: Our method chooses the shortest path between start node and end node in VG as the main stem, and each node on this stem is a natural hub of a community, as described below.

Let nodes 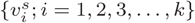 be the *k* nodes on the shortest path of start node and end node in a VG. 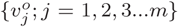 are the nodes outside the shortest path, where *m* = *N* − *k*. For any nodes 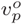 in 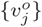, we compare the shortest path length between 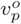 and each node in 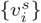, to identify the minimal path length value *l*_*pq*_ and corresponding node 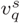 in 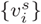. 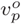 is then assigned to the community whose hub is 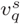. If there are more than one hub corresponding to the minimal value, we always choose the “left” hub, which corresponds to the earlier time point, as the target community’s hub. We then iterate through all nodes in 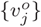, to get the initial community structure.

Finally, we measure the shortest path length between any pairs of hubs, i.e. the nodes in 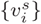, and if the shortest path length between them is less than a chosen cutoff, *є*, we then combine the two communities to obtain the final community structure.

Normally, the minimal *є* is the cutoff for which the network has the same number of communities as the number of hubs. Similarly, the maximal value of *є* corresponds to the case where the whole network becomes a single community. By changing the value of *є* between minimum and maximum, we can get the most stable community structure, i.e. the number of communities stabilizes when iterating with increasingly larger *є*. Thus, we can optimize the community structure. In our algorithm, we have provided the following options for cutoff selection: (i) with or without cutoff, (ii) fixed cutoff, or (iii) automatically optimized cutoff based on a plateau in number of communities as a function of *є*. This feature is unique to our method.

Another unique feature of our method is that we can choose the direction of how the nodes are connected in the community construction. Specifically, we can restrict the node 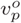 in 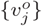 to only link to the community with hub 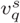 for which the corresponding time point *t*_*q*_ is earlier than the time point *t*_*p*_ corresponding to 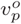. This feature essentially imposes a causality condition, where time points only depend on other past time points, and not future ones. It is important to keep time order in the community detection for characterizing biological time series from living systems. We have allowed flexibility in the implementation of the method, so the user can also choose node linking directions as earlier side, later side or both sides - this may be required for systems where there is time reversal symmetry.

Note that our method of shortest path based community detection may be used not only for WDPVG, but also for all kinds of visibility graphs, such as HVG and NVG.

Figure 1 provides a simple illustration of how our method works. A simulated time series is created from a cosine wave signal mixed with 40 percent random noise (Figure 1 A). Then, we construct the weighted NVG and the reflected perspective NVG in Figure 1 B and C. Afterwards, the weighted dual natural visibility graph is created by combining the NVG and the reflected perspective NVG (Figure 1 D). We use our community detection method to the graph in Figure 1 E showing the communities corresponding to the original time series.

**Fig. 1.**
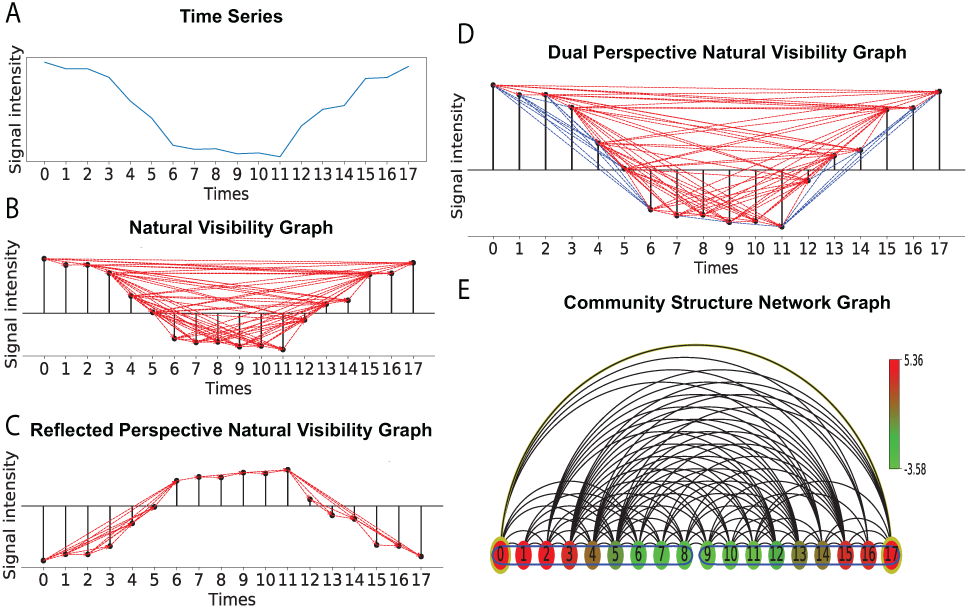
Illustration of the construction of weighted perspective visibility graph and community structure based on the shortest path length community detection method. **A** is the simulated time series cosine signal with 40 percent noise in intensity. We construct the weighted NVG and reflected NVG, where the edge weights are based on Euclidean distance. Edge connections of NVG and reflected NVG are illustrated in **B** and **C** respectively. The dual weighted perspective visibility graph is created by combining NVG and reflected NVG, as shown in **D**, where the links with red color come from NVG, and the blue links come from the reflected NVG. Using our shortest path length based community detection method, we can find the community structure of the time series, as shown in **E**. The time points in the same community are encircled by a blue outline, and the color of the nodes represents the signal intensity. The time series separates into two communities that capture the cosine signal’s periodicity.

### 2.3 Simulation

We used simulated time series to evaluate our method. We illustrate here three types of time series: cosine, square, and saw-tooth wave signals. Then we added different intensity random noise to each of these signals to test the tolerance of the community detection to noise. We also randomly removed different percentages of time points from these time series to check the robustness to missing data and the resulting uneven sampling. We then built the WDPVG for these time series, and detected communities using our method. The results of our method were compared to traditional community detection methods such as the Louvain method, which is a widely used method of fast greedy optimization of modularity (Blondel *et al*., 2008), and the Girvan–Newman hierarchical method which is based on centrality notions (Girvan and Newman, 2002). The Louvain method was implemented using the Python python-louvain package, (https://github.com/taynaud/python-louvain). The Girvan-Newman method is available in NetworkX, which is the most popular open source network analysis package in Python (Hagberg *et al*., 2008).

### 2.4 Experimental biological time-series applications

We also compared the results of our method to the Louvain and Girvan-Newman methods when applied to several experimentally acquired biological datasets. These datasets used are summarized below.

#### Saliva Set (DS1)

We used a saliva RNA-sequencing dataset we generated, that was obtained from a clinical trial monitoring individualized response to the standard 23-valent pneumococcal polysaccharide vaccinate (PPSV23). The saliva was sampled from a healthy individual. We had first carried out a 24-hour hourly sampling to establish a normal physiological state baseline. Then, we repeated with another 24-hour hourly sampling that also included vaccination with pneumococcal vaccine (PPSV23) to assess response to the vaccine. The vaccine was administered approximately 3.5 hours following the first hourly sample. Approximately 7 hours after vaccination, the individual reported having a fever that lasted about 4 hours. Here, we analyzed the differences between the two 24 hour periods: (i) the first 24 hours hourly sampling (Sal_1_(*t*)) and (ii) the 24 hours hourly sampling that included vaccination (Sal_2_(*t*)). We then constructed the paired difference time series Δ, where for each timepoint *i* for each gene *α*, Δ_*α*Sal_(*t*_*i*_) = Sal_2*α*_(*t*_*i*_) − Sal_1*α*_(*t*_*i*_). We carried out a categorization into groups and subgroups of gene expression based on these data (see online Python notebook, and previous discussion using PyIOmica (Domanskyi *et al*., 2019; Mias *et al*., 2016; Mias and Zheng, 2020)). For a given subgroup of genes, we constructed the mean time series across the members of this subgroup. We then built the WDPVG and compared the different temporal community detection methods results on this time series (see also online Python notebook for code and data at https://doi.org/10.5281/zenodo.3693984).

#### Diabetes Set (DS2)

The second dataset came from personal multi-omics profiling data (e.g. including blood measurements of A1C, fasting glucose and selected immune cytokines etc.) from individuals with Type 2 diabetes mellitus at its earliest stage (Zhou *et al*., 2019). As an example, we chose one individual’s A1C, fasting glucose and selected immune cytokines data from the rich dataset, as reported by the authors. These time series include 14 time points with different healthy condition. We constructed the VG from the time series and detected the communities, to assess whether our method can capture the physiological status of the subject for these time series.

#### Radiation Exposure Set (DS3)

Finally, we analyzed a radiation exposure time series dataset from wearable biosensors (Li *et al*., 2017). The data collected were hourly personal radiation exposure, assessed by a wearable biosensor for more than 100 days. We chose one day spans (24 hours, from 12 am to 11 pm) as the natural time window. We then analyzed separately four days when the individual of this study had flight activity, and the radiation reported on these days was higher than non-flight days. We then applied our methods to assess if we can detect the radiation events as community structures.

## 3 Results

### 3.1 Simulation

To investigate whether our method captures periodic features we simulated well defined periodic time series. We compared our path-length based method (PL) with two widely used community detection method, the Louvain method (LN) (Blondel *et al*., 2008) and the Girvan–Newman method (GN) (Girvan and Newman, 2002). Figure 2 shows the signal intensity and communities for the cosine and square wave signals’ time series (top and bottom, respectively). The communities our method detected matched exactly with the signal periods. The Louvain method also captured precise periodic features in the square wave signal, where it assigned communities corresponding to half periods. However, the Louvain method obtained some unmatched results in the case of the cosine signal time series. Finally, the Girvan–Newman method obtained coarser results compared to the other two methods.

**Fig. 2.**
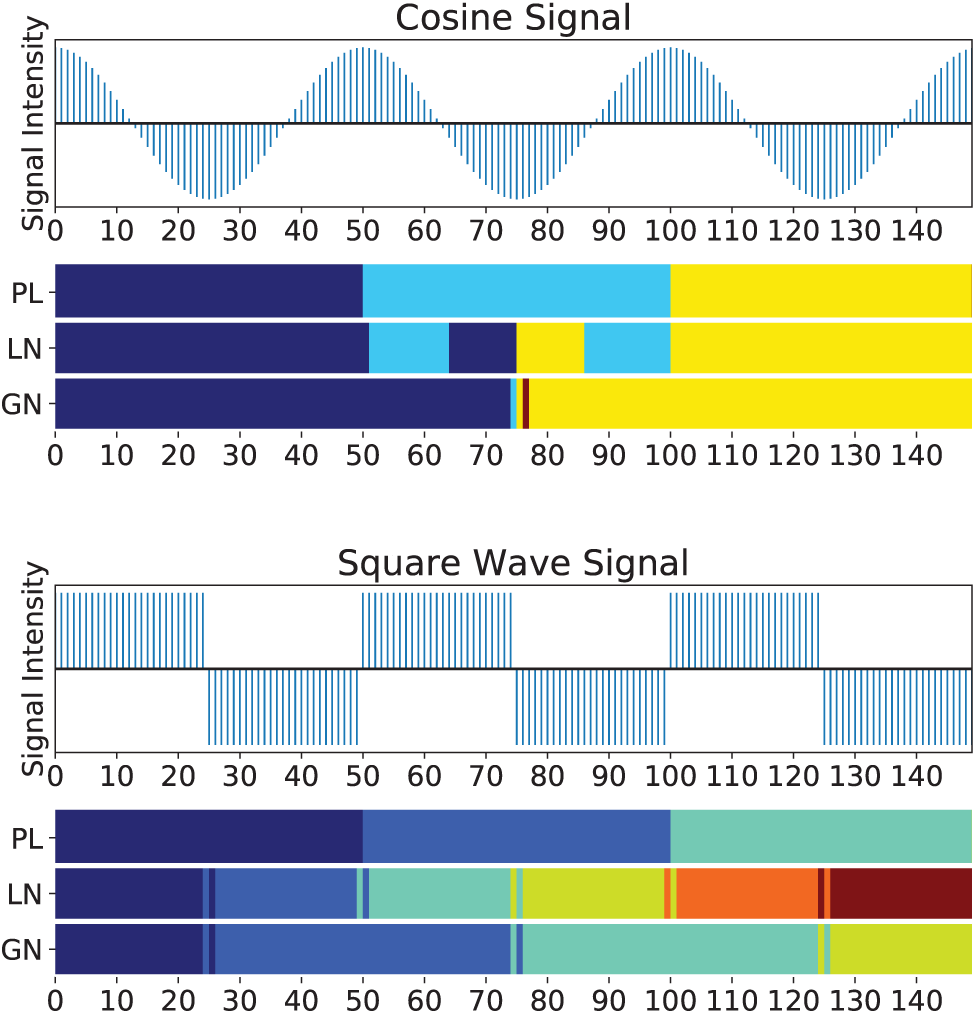
The intensity and communities of cosine (top) and square wave (bottom) signal time series. The communities obtained by our method (PL), Louvain method (LN) and Girvan–Newman method (GN) are presented with color bars, with time points in same community having the same color. The communities obtained with our method on both cosine and square wave time series are capturing the signal periods. The Louvain method works better on the square wave signal but worse on the cosine signal. The Girvan-Newman method captures the periods of the two time series with some errors at the boundaries.

In addition, Figure 3 shows results on the tolerance to noise and missing data for cosine wave (Figure 2A-D), square wave (Figure 2E-H), and saw-tooth wave signals(Figure 2I-L). Compared to the Louvain and Girvan–Newman methods, our method displays higher tolerance in either situation of noisy signals or uneven sampling. Whenever we added noise from 20 percent to 80 percent, or additionally removed time points from 10 percent to 40 percent, our method still captured the periodic changes. To the contrary, the results of the two traditional methods were irregular, with coarsely defined communities and multiple nodes in communities unmatched with the corresponding signal’s period. Even though the Louvain method worked well in the perfect square wave signal time series (e.g. without noisy and uneven sampling), the method showed low tolerance to noise or time point removal.

**Fig. 3.**
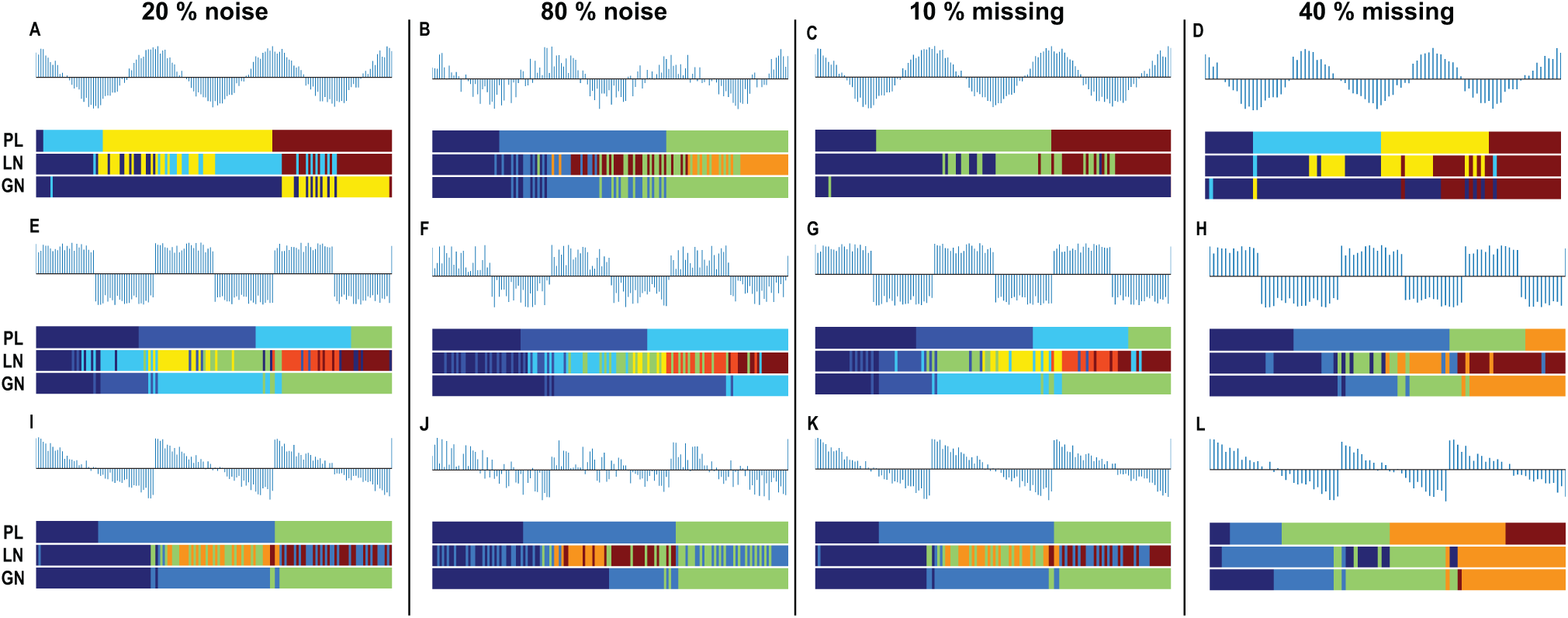
Community structure of different simulated signal time series. We simulated 3 types of signals: cosine time series **A-D**, square wave time series **E-H** and saw-tooth wave time series **I-L**. The community structure is represented by color bar, with time points in same community assigned the same color. For each kind of signal, we also added 20 percent and 80 percent noise (the first and second subplots of each row respectively) to test the tolerance to noise. We also randomly removed 10 percent and 40 percent time points (the third and forth subplots of each row respectively) to check the robustness to missing data and uneven sampling. We compared our method (PL, first color bar of each subplot) with the two traditional community detection methods, Louvain method (LN, second color bar) and Girvan–Newman method (GN, third color bar). Our method identifies community structures matching with the characteristics of the various time series, even in the presence of noise or missing data.

### 3.2 Experimental biological time-series applications

We then applied our method to the experimentally acquired biological time series summarized in the method section above. First we used our method to detect communities from the saliva omics monitoring experiment, DS1. We chose genes signals from the data displaying autocorrelation at lag 1 (simulation adjusted p-value < 0.01), and calculated the average across the signals for each time point. The average signal intensity is shown in Figure 4 A top. We then built the WDPVG from this signal and we obtained temporal communities using the different methods in Figure 4 A bottom. The community structure was found to reflect the four physiological states over the 24 hours: (i) pre-vaccination baseline, (ii) post-vaccination to fever onset, (iii) fever onset to resolution, (iv) post vaccination baseline. The PL method showed better separation of the periods corresponding to the physiological state of the subject. The LN and GN methods also performed well, displaying, however, some mixing of timepoints.

**Fig. 4.**
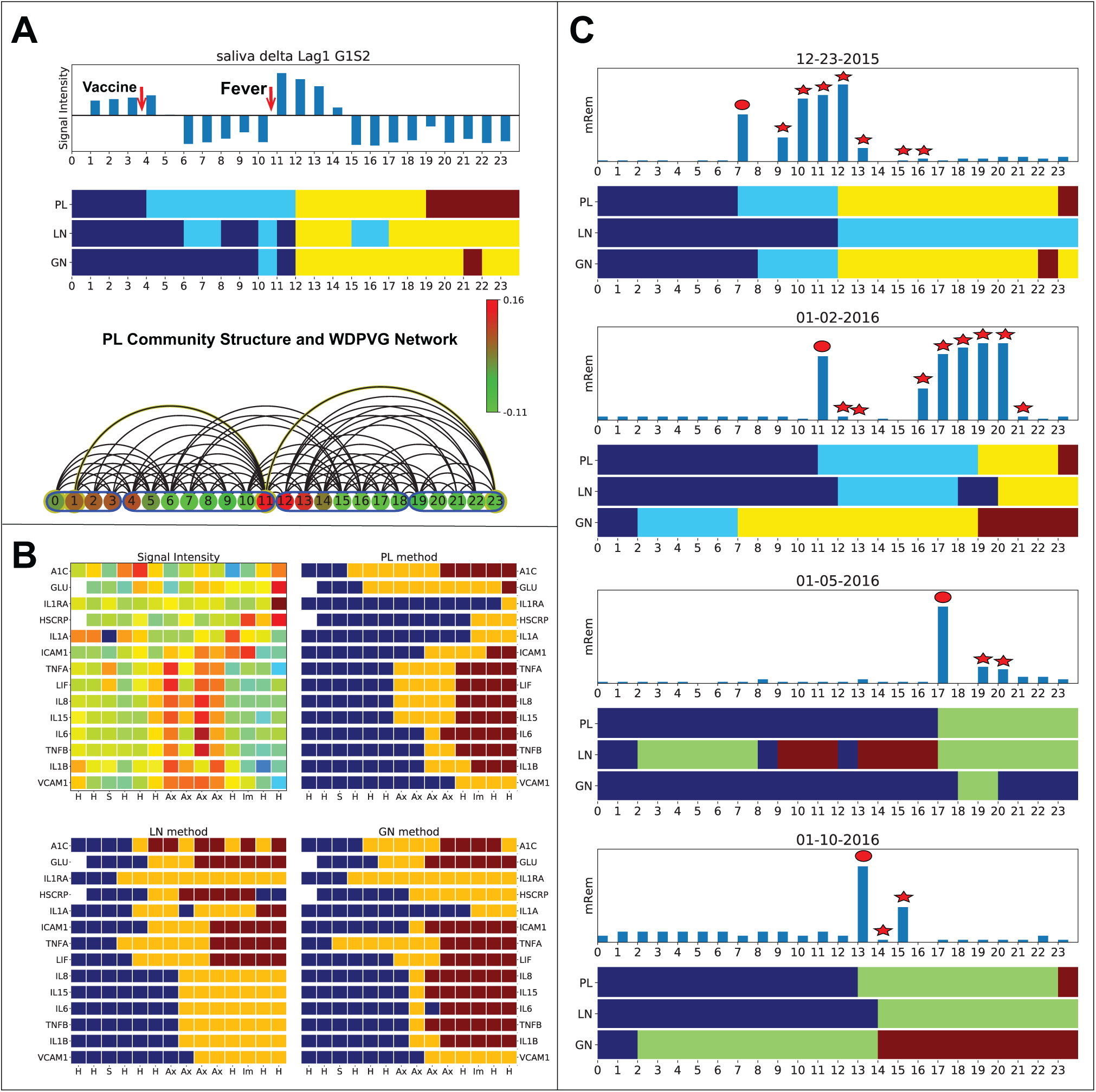
Results of our visibility graph based community detection method applied to experimental biological datasets. **A** shows the results of applying different methods to an individual’s saliva omics time series before and after their vaccination, over a 24 hour period. The time series indicates average gene expression for a subset of genes with autocorrelated behavior. The VG nodes’ color in **A** bottom represents the intensities of the time series, and nodes belonging to the same detected community are framed by a blue border. The communities correspond well to the four physiological subject states of biological significance: prior to vaccination, after vaccination, a post vaccination fever period and resolution, post fever relative recovery. **B** The standardized intensities of A1C, fasting glucose and selected immune cytokines time series from a subject with Type 2 diabetes (**B** top left) and their community structure (**B**left) are shown for different methods. The data are ordered temporally from left to right, and physiological states are indicated per timepoint as one of H (healthy), S (stress), Ax (antibiotic regiment) and Im (immunization) states. For the experimental data, warm colors indicate higher and colder colors lower intensities respectively. In the methods’ results, nodes belonging to the same community are depicted with the same color bar. The communities structure reflects the changes in physiological state that result from molecular intensity differences. **C** The radiation intensity data from wearable biosensors in four separate days including flight activity are shown. The red disc indicates that the radiation was recorded during the airport carry-on luggage check. Stars represent radiation monitored during flight timepoints. Again, nodes in the same community are indicated with the same color in the horizontal bars. The PL community structures during each of these four days indicate the periods of varying radiation exposure, and correctly identify the onset of the exposure.

The analysis of the Type 2 diabetes dataset, DS2, focused on results previously reported (see Figure 6B in Zhou *et al*. (2019)). The data included signals of A1C, fasting glucose and selected immune cytokines. There are no negative values in the dataset, so we built a normal perspective NVGs from each entry, and obtained the community structures. Figure 4 B top left shows the heatmap of standardized reported intensities (i.e. the results of (Zhou *et al*., 2019), with the communities structures heatmaps, shown for PL, LN and GN methods as well in Figure 4 B respectively. The community structures detected for DS2 reflect the change of time series intensities changes. We note that our community detection method does not require standardization of the raw data. From the figure we see that the community changes capture the status changes in each signal, while effectively filtering out the noise in the data. The PL method results overall better reflect the duration of the different physiological states, as compared to the LN and GN methods.

Finally, we applied our method to four separately days’ radiation exposure measurements from DS3. We show the four separate day results with high radiation exposure changes (when the individual was traveling by flight on these days) in Figure 4 C. The community structures of these four days all capture the radiation exposure changes, acting as an adverse event detector. Again, the PL method is more concordant with the radiation exposure timeframes overall, without mixing of timepoint in the communities.

## 4 Conclusion

We have introduced new methods to characterize graphs derived from time series through the application of VGs. We have introduced WDPVGs that combine normal perspective and reflected perspective visibility graphs, so that the peaks and troughs of a time series can be simultaneously represented. The WDPVGs also take into account uneven sampling effects through weight assignments to the edges. The WDPVG approach thus produces a graph that captures well the characteristics of the underlying time series.

We have also developed a new method to detect communities of VG. Our VG community detection method is based on the graphs geometry and considers the shortest path from the start node to the end node. The method does not assume a random graph null model. This makes the method advantageous and more appropriate for all kinds of VGs (e.g. NVG, HVG or WDPVG), because VGs cannot be compared to random graphs due to the sequential nature of time points.

The several simulated time series we used to test our WDPVG and VG community detection methods supported their validity. Our methods also showed high tolerance for uneven sampling and signal noise. Our PL community detection method showed robustness to noise compared to traditional community detection methods such as the Louvain and Girvan–Newman methods. Overall, the results suggest that our approach is better suited for community detection within VGs.

The application of our method to experimental biological datasets gave examples of how the method may be used to identify temporal communities that correspond to biological states (e.g. physiological state of a subject, changes in molecular measurements due to vaccination, or detection of radiation exposure). The method has great potential not only for detecting the boundaries of biological temporal states, but for medical implementation in detecting potential adverse medical events from temporal measurements.

## Funding

This work was supported by the Translational Research Institute for Space Health through NASA Cooperative Agreement NNX16AO69A (project T0412). S.D. acknowledges support by the NIH under R01GM122085.

## Notes

https://doi.org/10.5281/zenodo.3693984

